# Tissue-Specific Activation of Microhomology-Mediated End Joining with Age Reflects Dynamic Rewiring of DNA Repair in Rats

**DOI:** 10.1101/2025.06.13.659285

**Authors:** Diksha Rathore

**Author notes:** **To whom correspondence should be addressed:** Department of Biochemistry, Indian Institute of Science, Bangalore-560012, India.

## Abstract

Genomic instability is a hallmark of ageing, driven by the accumulation of DNA lesions and the decline of high-fidelity repair pathways such as homologous recombination (HR) and classical nonhomologous end joining (c-NHEJ). We hypothesized that ageing cells increasingly rely on microhomology-mediated end joining (MMEJ), an inherently error-prone repair pathway, whose role in ageing remains poorly characterized. Here, we present the first systematic, tissue-wide analysis of MMEJ activity across eight rat organs using an *in vitro* assay with defined 10–16 nt microhomology substrates and cell-free extracts from rats of different ages. Our results reveal age-associated and tissue-specific reprogramming of MMEJ: activity increased in the testes and liver, newly emerged in kidneys and lungs, and declined in immune tissues such as the spleen and thymus. Strikingly, aged brain extracts exhibited latent MMEJ activity. Repair efficiency was further modulated by microhomology length in a tissue-dependent manner. These functional shifts correlated with altered expression of key MMEJ effectors, including XRCC1, PARP1, Ligase III, and notably FEN1, whose upregulation in aged lungs mirrored robust MMEJ activation. Together, our findings uncover dynamic remodelling of MMEJ during ageing, implicating it in age-related genomic instability and highlighting potential therapeutic targets to mitigate mutation burden in ageing tissues.

## Introduction

Ageing is characterized by a decline in cellular function and a reduced DNA repair capacity, leading to DNA damage accumulation. This damage builds up over time due to prolonged exposure to damaging agents, contributing to the development of age-related diseases (1). This diminished repair manifests as elevated somatic mutation rates, which, however, differ in extent and type across tissues. For example, ageing lymphocytes display a pronounced rise in mutations at the hypoxanthine-guanine phosphoribosyl transferase (HPRT) locus (2,3). In murine models, the bladder and liver accumulate mutations steadily with advancing age, whereas the brain shows no net increase in mutations after 4-6 months (4,5).

Additionally, distinct mutations become prevalent with age in proliferating versus post-mitotic tissues. Specifically, point mutations are more common in the small intestine, while genomic rearrangements are predominantly observed in the heart (6). These findings indicate that DNA damage and mutations increase with age. Among the various types of mutations, genomic rearrangements appear to have a particularly prominent impact on ageing (7,8)These rearrangements primarily arise from erroneous repair of DNA DSBs, which are known to accumulate with age(9). This accumulation, along with age-associated shifts in DSB repair pathway usage, supports the idea that impaired DSB repair is a key driver of the ageing process (9–14).

In parallel, ageing also impairs the repair of point mutations, which are repaired by Mismatch repair (MMR), base excision repair (BER), and nucleotide excision repair (NER)(15–18). In ageing cells, MMR activity decreases, as shown in human T-cell clones with increased passage numbers (19). BER also declines, resulting in the accumulation of oxidative damage and AP sites, particularly in tissues like the liver and heart (20), and reduced repair enzyme activity (21). NER, responsible for repairing UV-induced damage, studies showed age-related decline, with reduced repair efficiency observed in human fibroblasts and rodent models (22,23).

Beyond point mutations, ageing cells must also contend with the repair of DSBs, which pose a severe threat to genome integrity (9). DSBs are repaired through various pathways, including Homologous Recombination (HR), Nonhomologous End Joining (NHEJ), Microhomology-Mediated End Joining (MMEJ), and Single-Strand Annealing (SSA). NHEJ is the primary repair mechanism in mammalian cells; it is fast but can introduce small deletions and insertions (indels) (24). HR, MMEJ, and SSA rely on homologous sequences for repair, with HR being highly accurate. However, both SSA and MMEJ tend to cause deletions of intervening sequences between repeats, which can lead to genomic rearrangements and mutations (25,26).

With age, the efficiency of DSB repair mechanisms, particularly NHEJ, declines significantly. This age-related decline has been documented in multiple tissues. For instance, studies on rats revealed significant reductions in NHEJ activity in older brains and neurons (27,28) this decline is often attributed to a decrease in Ku70 and DNA-dependent protein kinase (DNA-PK) activity, a key component of the NHEJ machinery (29). Similar findings show impaired NHEJ activity in Alzheimer’s patients (30). In senescent cells, primary components of NHEJ, such as Ku70 and Ku80, are decreased, and their distribution within the nucleus and cytoplasm becomes disrupted. This results in impaired DNA repair (31) (32). In human studies, a decline in DSB repair efficiency with ageing has been confirmed, including a decrease in the γ-H2AX response and reduced expression of core NHEJ components such as XRCC4 and Lig4 (33,34). Age-related changes in repair mechanisms are also observed in human lymphocytes, where levels of Ku70 and MRE11 decrease with age, further supporting the idea of compromised DNA repair with age (35). Additionally, a reduction in BRCA1 and ATM-mediated repair in ovarian follicles contributes to increased DSBs in ageing mice and women with BRCA1 mutations (10,36).

Changes in HR efficiency with age contribute to genomic instability and increased disease susceptibility. Some studies show a decline in HR efficiency with ageing, such as in Drosophila and human cells, where Rad51 recruitment increases but repair remains less effective (37,38). In *Drosophila*, aged male germ cells appear to compensate for diminished NHEJ efficiency by upregulating HR (39). However, this compensatory shift may paradoxically exacerbate genomic instability. These observations highlight the conflicting evidence surrounding age-associated changes in HR efficiency.

This complexity is mirrored in other alternative repair pathways. For example, while some studies report a decline in SSA efficiency with age (40). Others have observed increased SSA activity, particularly in the context of breast cancer, where elevated SSA is likely associated with heightened genomic instability and tumorigenesis (41).

Importantly, a consistent trend observed across species and tissues is the age-associated shift from high-fidelity repair mechanisms toward more error-prone alternatives. In aged mice, there is a significant decline in the efficiency and fidelity of cNHEJ, accompanied by an increase in MMEJ events, particularly within the lungs and heart (42). Similarly, in *Schizosaccharomyces pombe*, ageing drives genome-wide rearrangements characterized by microhomology at breakpoint junctions, indicative of MMEJ activity(43). This age-associated shift towards MMEJ is also observed in aged breast cancer cells (41), suggesting a conserved mechanism of genomic instability across species.

Although age-associated declines in canonical DNA repair pathways such as HR and cNHEJ are well characterized, evidence for a compensatory increase in error-prone pathways, particularly MMEJ, remains limited, and the mechanistic basis of this shift is not yet understood. Additionally, molecular drivers responsible for the age-associated changes in MMEJ have yet to be identified. Given the inherently mutagenic nature of MMEJ and its potential contribution to age-related genomic instability and disease, including cancer and degenerative disorders, this knowledge gap represents a critical frontier in ageing and genome maintenance research.

Targeted investigation into the regulation of MMEJ with age could uncover novel therapeutic opportunities to mitigate the deleterious consequences of genomic instability in aged tissues. Defining the protein drivers of this age-associated shift is crucial for restoring repair fidelity and maintaining genome integrity in ageing tissues.

Here, we present a comprehensive investigation into how ageing modulates MMEJ activity across multiple mammalian tissues, using a defined *in vitro* end-joining assay with substrates bearing controlled microhomology lengths. By profiling MMEJ efficiency in eight distinct rat organs across three physiologically relevant age groups, we uncover a dynamic and tissue-specific reprogramming of this error-prone repair pathway. These functional changes correlate with differential expressions of key MMEJ factors, revealing potential molecular drivers of repair plasticity during ageing. Together, our study provides mechanistic insight into the shifting landscape of DNA repair fidelity with age and identifies MMEJ as a previously underappreciated contributor to age-associated genomic instability. Importantly, these findings open new avenues for therapeutic intervention, suggesting that selective modulation of MMEJ components may mitigate mutagenic repair and reduce the burden of age-related diseases driven by genome instability

## Experimental procedure

### Enzymes, chemicals, and reagents

The chemicals and reagents used in this study were obtained from Amresco (USA), SRL (India), Sigma Chemical Co. (USA), and Himedia (India). Restriction enzymes and other DNA-modifying enzymes were sourced from New England Biolabs (USA) and Fermentas (USA). Radiolabeled nucleotides were provided by BRIT (India) and Revvity (USA). Antibodies were purchased from BD (USA), Cell Signalling Technology (USA), Santa Cruz Biotechnology (USA), and Calbiochem (USA). Oligomeric DNA was procured from Eurofins Genomics (India), Juniper LifeSciences (India), Medauxin (India), and Xcelris Genomics (India).

### Oligomeric DNA

The oligomers used in this study are detailed in Tables 1 and 2. These oligomeric DNA were purified using 12-15% denaturing PAGE whenever needed. The relevant section on materials and methods provides specific information regarding the use of oligomers in each experiment. For clarity, a simplified nomenclature is applied to some of these oligomers when presenting experimental results.

**Table1.**
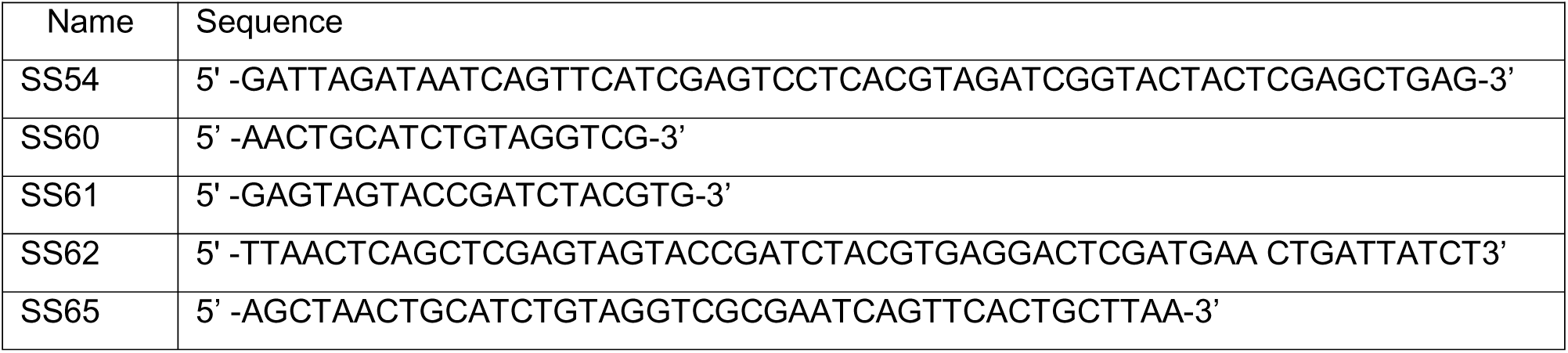
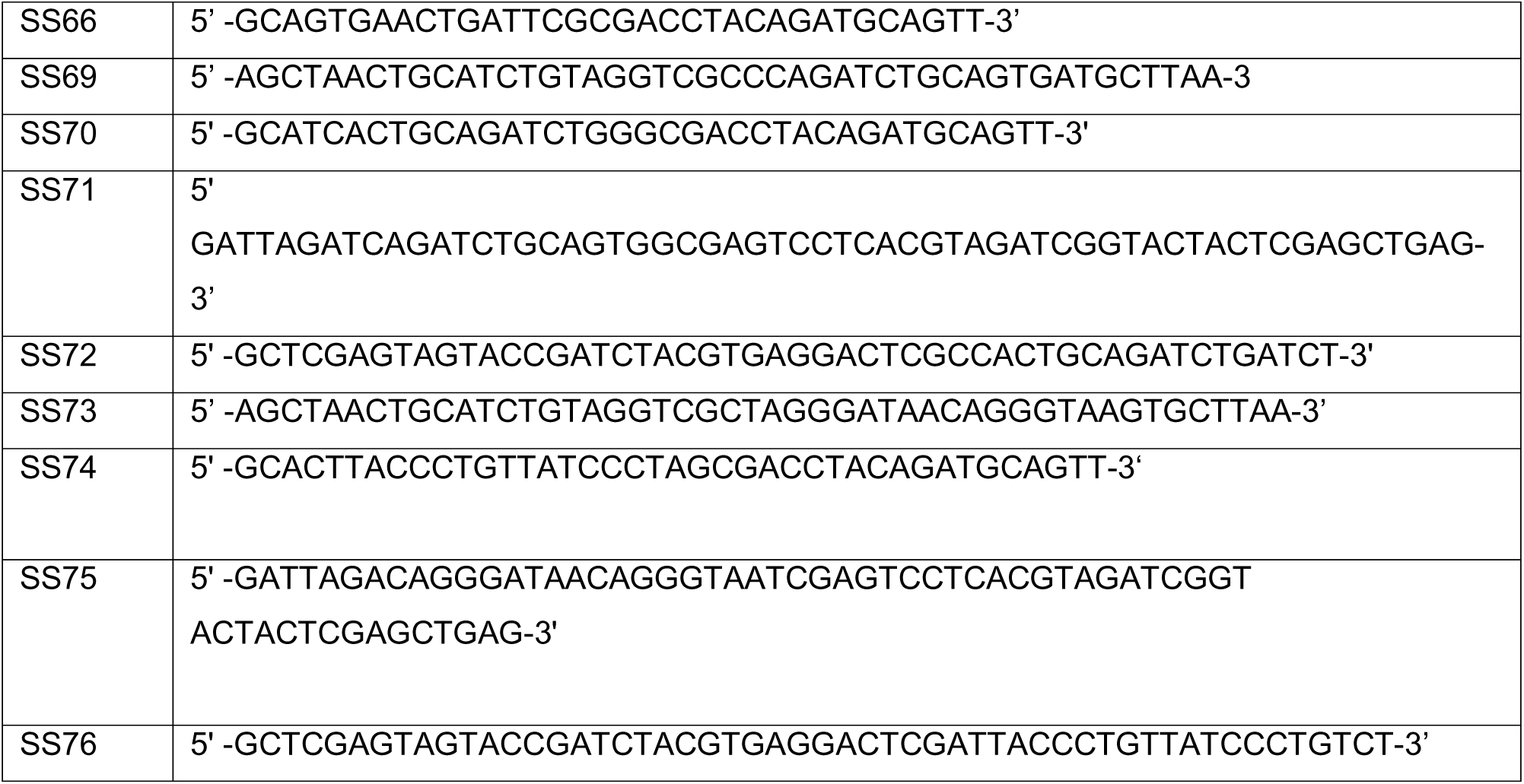

**Table 2.**

| Microhomology length | Substrate I | Substrate II |
| --- | --- | --- |
| 10 | SS54/SS62 | SS65/SS66 |
| 13 | SS69/SS70 | SS71/SS72 |
| 16 | SS72/SS73 | SS74/SS75 |

### Preparation of dsDNA substrates from oligomers

Double-stranded DNA substrates mimicking breaks flanked by direct repeats were generated by annealing complementary oligonucleotides in a buffer containing 100 mM NaCl and 1 mM EDTA, as described previously (44). The study used microhomology regions of varying lengths: 10, 13, and 16 nucleotides (see Tables 1 and 2). As described earlier, the 5′ end-labelling of SS60 and other oligomers with γ^−32^P ATP was performed using T4 polynucleotide kinase, and the labelled oligomers were stored at −20°C.

### Animals

Male Wistar rats (*Rattus norvegicus*), aged 1–2 months (young), 3–4 months (middle-aged), and 6–8 months (aged), were obtained from the Central Animal Facility of the Indian Institute of Science (IISc), Bangalore, India. All procedures were conducted per the Institutional Animal Ethics Committee guidelines (CAF/Ethics/026/2023) and adhered to national regulations for the care and use of laboratory animals.

Animals were housed in a controlled environment with regulated temperature and humidity and maintained on a 12 h light/dark cycle. Rats were housed in standard polypropylene cages, providing ad libitum water access and a nutritionally balanced pellet diet (Agro Corporation Pvt. Ltd., India). The diet comprised 55% nitrogen-free extract (carbohydrates), 21% protein, 5% lipids, 4% crude fibre, 8% ash, 1% calcium, 0.6% phosphorus, 3.4% glucose, and 2% vitamins.

### 5’ end labelling of oligomeric substrates

The 5’ end of the oligomeric substrates was labelled using γ-³²P ATP and 1 unit of T4 polynucleotide kinase in a reaction buffer composed of 20 mM Tris-acetate (pH 7.9), 10 mM magnesium acetate, 50 mM potassium acetate, and 1 mM DTT. The reaction was carried out at 37°C for 1 h. Following labelling, the radiolabeled oligomers were purified using a Sephadex G25 column and stored at −20°C (45,46).

### Preparation of cell-free extracts from various rat organs

Cell-free extracts were generated from various organs of male Wistar rats aged 1–2 months, 3–4 months, and 6–8 months, including the testes, brain, lungs, heart, spleen, kidneys, liver, and thymus, following the protocol outlined by (Baumann & West,1998), (47)with slight modifications. Initially, the tissues were rinsed with PBS and washed in a hypotonic buffer (10 mM Tris-HCl, pH 8.0; 1 mM EDTA; 5 mM DTT) by centrifugation at 2000 rpm for 10 min at 4°C. The tissue pellets were then resuspended in twice their volume of the same hypotonic buffer and left on ice for 20 min before homogenization. A cocktail of protease inhibitors, including 0.01 M phenylmethylsulfonyl fluoride, 2 μg/ml aprotinin, 1 μg/ml pepstatin, and 1 μg/ml leupeptin, was added to prevent protein degradation. After incubating the homogenate on ice for another 20 min, a high-salt buffer (50 mM Tris-HCl, pH 7.5; 1 M KCl; 2 mM EDTA; 2 mM DTT) was added at a volume equivalent to half of the homogenate. The mixture was then centrifuged at 42,000 rpm for 3 h at 4°C using a Beckman TLA-100 rotor. The resulting supernatant was dialyzed overnight at 4°C against a dialysis buffer (20 mM Tris-HCl, pH 8.0; 0.1 M potassium acetate; 20% glycerol; 0.5 mM EDTA; 1 mM DTT). After dialysis, the extracts were snap-frozen and stored at −80°C. Protein concentrations were determined using the Bradford assay, and equal loading was confirmed by SDS-PAGE followed by Coomassie Brilliant Blue staining.

### MMEJ assay

MMEJ activity was assessed as previously described (45,46,48). With minor modification, briefly, DNA substrates containing microhomology regions of varying lengths (10, 13, and 16 nucleotides) were incubated with cell-free extracts in a reaction buffer containing 50 mM Tris-HCl (pH 7.6), 20 mM MgCl₂, 1 mM DTT, 1 mM ATP, and 10% polyethylene glycol (PEG). Reactions were carried out at 30°C or 37°C for the indicated time periods, depending on the experimental conditions.

Following incubation, the reactions were terminated by heating at 65°C for 20 min to denature proteins. The end-joined DNA products were amplified by PCR using a radiolabeled forward primer (SS60) and an unlabeled reverse primer (SS61) under the following thermal cycling conditions: an initial denaturation at 95°C for 3 min, followed by 15 cycles of 95°C for 30 s, 58°C for 30 s, and 72°C for 30 s, with a final extension step at 72°C for 3 min.

Amplified products were resolved on a 10% denaturing polyacrylamide gel. A 60-nucleotide radiolabeled oligonucleotide was included in each gel as a size marker, with the expected MMEJ product being 62 nucleotides. Gels were imaged using a phosphorimager (Fuji, Japan), and band intensities were quantified using Multi Gauge software (Version 3.0).

### Quantification

We used Multi Gauge (V3.0) software to measure the bands of interest. We selected a rectangle around each band to measure its intensity and did the same for all bands in each lane. We also measured the background intensity from a blank area of the same lane and subtracted it. The final intensity values were plotted and shown in a bar graph.

### Statistical analyses

Values were presented as mean ±S.E.M. for both control and experimental samples. Statistical analyses were conducted using one-way ANOVA followed by an unpaired Student’s t-test with GraphPad Prism 6 software (San Diego, CA, USA.). Results were considered statistically significant if the p-value was ≤0.05.

## Results

### MMEJ is preferentially active in proliferative tissues (thymus, spleen, testes, and liver) of young rats

Multiple lines of evidence suggest that the error-prone repair pathway, MMEJ, plays a critical role during mitosis (49–51). Supporting this, our unpublished data (Rathore, 2025) reveal that MMEJ efficiency varies significantly across different tissues in young rats. Higher levels of repair activity were observed in highly proliferative tissues such as the thymus, spleen, testes, and liver. In contrast, markedly lower activity was detected in non-dividing tissues, including the brain, lungs, heart, and kidneys (Figure 1a–d). These findings align with previous studies reporting elevated MMEJ activity and increased expression of DNA polymerase theta (Polθ)—a crucial MMEJ enzyme—in proliferative tissues such as the testes and thymus (44,52–54). Building on our initial observations, the current study aimed to investigate how MMEJ repair capacity is modulated with age across multiple tissue types. Specifically, we assessed how this DNA repair pathway changes as rats transition from the young to middle-aged and aged stages of life.

**Figure 1:**
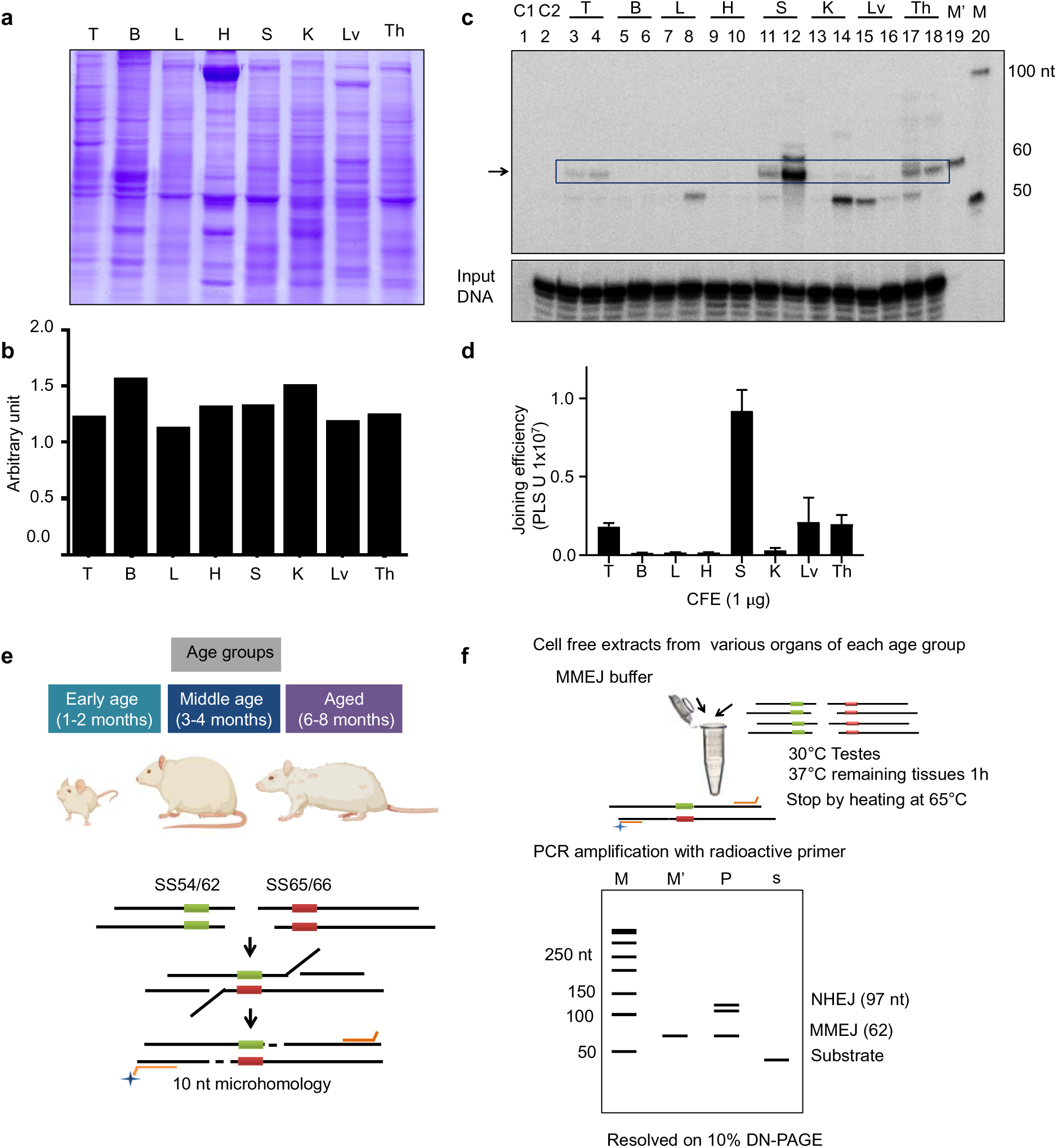
Tissue-specific variation in MMEJ efficiency in young rats. (**a**) SDS-PAGE profile of cell-free extracts (CFEs) prepared from various rat tissues: testis (T), brain (B), lungs (L), heart (H), spleen (S), kidneys (K), liver (Lv), and thymus (Th). (**b**) Bar graph representing protein normalization based on the gel profile shown in panel (a). (**c**) MMEJ activity across tissues was assessed by incubating 1 µg of each CFE with a DNA substrate mimicking a double-strand break flanked by 10-nt direct repeats. Reactions were carried out in MMEJ buffer (10 mM Tris-HCl, pH 8.0; 20 mM MgCl_₂_; 1 mM ATP; 10% PEG 8000; 1 mM DTT) for 1 hour at 30°C for testis extracts and 37°C for all other tissues. Joining products were resolved on 10% denaturing PAGE and visualized alongside a 50-nt DNA ladder (M). (**d**) Quantitative comparison of MMEJ efficiency from 1 µg of CFE derived from each tissue. Data represents the mean ± S.E.M. from three independent experiments using extracts from separate animals. (**e**) Schematic overview of the experimental design. MMEJ activity was evaluated across three age groups—young (1–2 months), middle-aged (3–4 months), and aged (6–8 months)—using substrates with 10-nt microhomology, bottom panel explain the repair mechanism via MMEJ (**f**) Diagram of the in vitro MMEJ assay workflow. Normalized CFEs from each age group were incubated with a microhomology-flanked substrate. Joined products were detected by radiolabeled PCR and resolved on 10%denaturing PAGE. ‘M’ indicates a 50-nt radioactive marker; ‘M’’ represents the 60-nt MMEJ-specific oligomer product; ‘S’ denotes the unprocessed substrate; ‘P’ means the product formed via MMEJ-mediated repair.

### Tissue and age-dependent Profiling of MMEJ activity in rats using defined microhomology substrates

To systematically investigate how MMEJ activity is changes across different tissues and stages of aging, we generated cell-free extracts (CFEs) from eight organs— testes, brain, lungs, heart, spleen, kidneys, liver, and thymus—isolated from male Wistar rats representing three age groups: young (1–2 months), middle-aged (3–4 months), and aged (6–8 months) (Figure 1e). CFEs were prepared following the protocol of Baumann and West (1998) (47). MMEJ activity was assessed using two synthetic DNA substrates (SS54/62 and SS65/66) harbouring a 10-nucleotide microhomology region flanking a defined double-strand break (Figure 1e). These substrates were incubated with CFEs, allowing endogenous repair proteins to mediate end joining. Reactions were carried out at 30 °C for testes extracts, to reflect their physiological conditions during spermatogenesis, and at 37 °C for all other tissues. The joined product (62 nt) was detected via radiolabeled PCR, and its intensity was used as a direct readout of MMEJ efficiency (Figure 1f).

### Middle age triggers the induction of MMEJ in the kidneys

Firstly, we analyzed changes in MMEJ activity during middle age by preparing CFEs from eight distinct organs of 3–4-month-old rats, representing the transition from youth to middle age. Total protein amounts were normalized using 8% SDS-PAGE to ensure equal protein input across tissues, enabling accurate comparison of MMEJ activity (Figure 2a, 2b). Repair reactions were carried out with both 1 µg and 2 µg of protein to assess dose-dependent differences in end-joining efficiency (Figure 2c, d, and e). As expected, no joined product was observed in reactions lacking protein input (Figure 2c, lane 2), confirming that the detected MMEJ activity was entirely protein-dependent and not due to spontaneous annealing or PCR artefacts.

**Figure 2:**
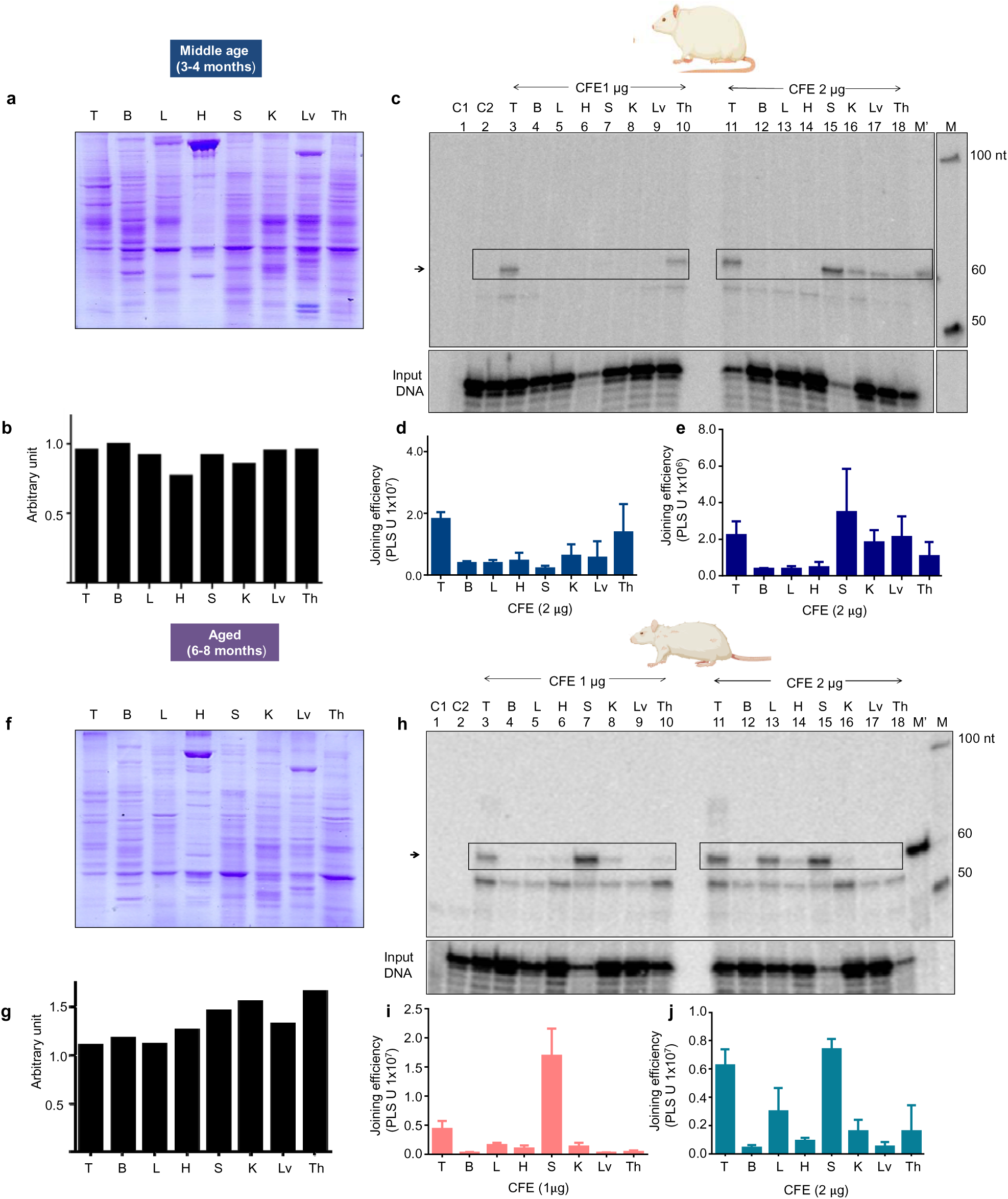
CFE Normalisation and assessment of MMEJ efficiency across various organs of middle-aged and aged rats. **(a)**SDS-PAGE analysis showing the normalization of CFE (Cell Free Extract) from various organs of middle-aged (3-4 months) rats, including testes (T), brain(B), lungs(L), heart (H), spleen(S), kidneys (K), liver (Lv), and thymus(Th). **(b)** A bar diagram showing the normalization profile for proteins in PAGE is shown in panel (c). A comparison of MMEJ efficiency across different organs in middle-aged rats is shown. A 10 nt microhomology substrate was incubated with 1 µg (lanes 3-10) or 2 µg (lanes 11-18) of CFE from the specified rat organs. The joining products were amplified using a radioactive primer and resolved on a 10% denaturing PAGE gel. M represents the 50 nt ladder, and M’ denotes the 60 nt marker. Lane 1 is the no-template control, and lane 2 is the no-protein control. **(d)** Bar graph showing the intensity of MMEJ joining products obtained with 1 µg of CFE from various organs of middle-aged rats. (**e)** Bar graph showing the intensity of MMEJ joining products obtained with 2 µg of CFE from various organs of middle-aged rats. **(f)** SDS-PAGE analysis showing the normalization of CFE from various organs of aged (6-8 months) rats **(g)** A bar diagram showing the normalization profile for proteins in PAGE is shown in the panel (**f**). **(h)** Comparison of MMEJ efficiency across different organs in aged rats. A 10 nt microhomology substrate was incubated with 1 µg (lanes 3-10) or 2 µg (lanes 11-18) of CFE from the specified rat organs. The joining products were amplified using a radioactive primer and resolved on a 10% denaturing PAGE gel. M represents the 50 nt ladder, and M’ denotes the 60 nt marker. Lane 1 is the no-template control, and lane 2 is the no-protein control. **(i)** Bar graph showing the intensity of MMEJ joining products obtained with 1 µg of CFE from various organs of late-aged rats. The results are from at least three independent experiments, with joining efficiency labelled in PSLU. Error bars represent the S.E.M. (j). Bar graph showing the intensity of MMEJ joining products obtained with 2 µg of CFE from various organs of late-aged rats. The results are from at least three independent experiments, with joining efficiency labelled in PSLU, and error bars represent the S.E.M.

At a protein concentration of 1 µg, the MMEJ activity profile in mid-aged rats partially resembled that of their younger counterparts. Repair efficiency remained highest in the testes and thymus—tissues known for their robust proliferative activity— while the brain, lungs, kidneys, heart, spleen and liver showed only minimal joining (Figure 2c, lanes 3–10). Quantification confirmed this trend, reinforcing the idea that highly proliferative tissues are privileged sites of MMEJ engagement (Figure 2d). Interestingly, at 2 µg protein concentration, subtle yet distinct changes in the repair profile became evident (Figure 2c lane 11-18). The spleen displayed intense joining at 2 µg CFE (Figure 2c, lane 15). In contrast, the kidneys, which were largely inactive in young animals (Figure 1c, 1d), began to exhibit emergent MMEJ activity at 2 µg (Figure 2c, lane 16). This activation of repair in previously quiescent or less responsive tissues suggests a dose-dependent engagement of the MMEJ pathway and may reflect a compensatory upregulation in response to accumulating DNA damage during mid-life. These results suggest that MMEJ is not static but undergoes age-dependent modulation.

In contrast, MMEJ efficiency in thymic extracts declined at the higher protein concentration (Figure 2c, lane 18). This reduction may signal a saturation effect or early degenerative changes in immune tissues, consistent with the well-documented thymic involution during ageing (55). Thus, middle age represents a pivot point in MMEJ regulation, characterized by both functional gains in certain tissues and early declines in others.

### MMEJ emerges in the lungs of aged rats

To understand how these trends evolve into late age, we extended our analysis to 6–8-month-old rats. CFEs from all eight tissues were normalized (Figure 2f, 2g), and the MMEJ assay was repeated at 1 µg and 2 µg protein (Figure 2h).

At a protein concentration of 1 µg, the MMEJ activity profile in aged rats exhibited a distinct shift relative to middle-aged counterparts. The testes continued to demonstrate robust repair, maintaining MMEJ efficiency well into old age, remarkable preservation given the age-related decline typically observed in DSB repair pathways (27,28,37). In contrast, the spleen, which exhibited minimal activity at this concentration in middle age groups, displayed a marked increase in MMEJ, while the kidneys demonstrated moderate repair activity similar to middle age. Conversely, thymic activity declined further, consistent with progressive impairment of repair capacity. These observations highlight a tissue-specific remodelling of MMEJ during ageing, suggesting both preservation and compensatory activation of repair mechanisms in select tissues (Figure 2h, 2i).

At a protein concentration of 2 µg, the MMEJ activity profile in aged animals partially resembled that of middle-aged rats, with high levels of end-joining observed in the testes and spleen. The most striking age-associated alteration was detected in the lungs, which exhibited a robust MMEJ activity (Figure 2h, lane 13)—a stark contrast to the negligible levels detected in younger (Figure 1c, 1d) and middle-aged rats (Figure 2c, 2e). This late-onset induction in a post-mitotic tissue suggests a potential compensatory response to persistent DNA damage or cellular stress. Meanwhile, the thymus continued to exhibit a further reduction in MMEJ activity, underscoring a sustained decline in repair proficiency. Collectively, these results support a model in which ageing induces a complex and tissue-specific reconfiguration of MMEJ capacity, encompassing both compensatory upregulation and progressive loss across different organ systems.

To ensure these observations reflected true biological differences and not technical variability, CFEs from all eight organs across the early, middle, and aged groups were standardized based on total protein content by resolving the extracts on an 8% SDS-PAGE gel (Supplementary Figure S1a–h). This approach controlled potential technical variability and ensured that observed changes in repair activity were not due to unequal CFE. For instance, testes CFEs were normalized across all age groups to confirm that differences in MMEJ activity arose from age-dependent biological changes rather than inconsistencies in CFE amounts.

### Tissue-specific age-dependent changes in MMEJ activity using a 10-nt microhomology substrate

Following normalisation, extracts were incubated with a DSB substrate containing a 10-nt microhomology, a well-characterised target for MMEJ. Incubation conditions were optimised for tissue-specific enzymatic activity: testis extracts were incubated at 30°C, while extracts from other organs were incubated at 37°C. This experimental framework enabled a controlled, reproducible comparison of MMEJ activity across tissues at different ageing stages, strengthening the validity of our findings on MMEJ repair dynamics in ageing. Our analysis revealed a striking, tissue-specific pattern of age-associated modulation in MMEJ activity. In testes, MMEJ efficiency remained stable from young to middle age but increased significantly in the aged group (Figure 3a, b; lanes 2–4), suggesting a late age enhancement of MMEJ in germline tissue. In sharp contrast, brain extracts exhibited a complete absence of MMEJ across all age groups (Figure 3a, c; lanes 5–7), underscoring a persistent suppression or lack of engagement of this error-prone pathway in non-dividing neural tissue.

**Figure 3:**
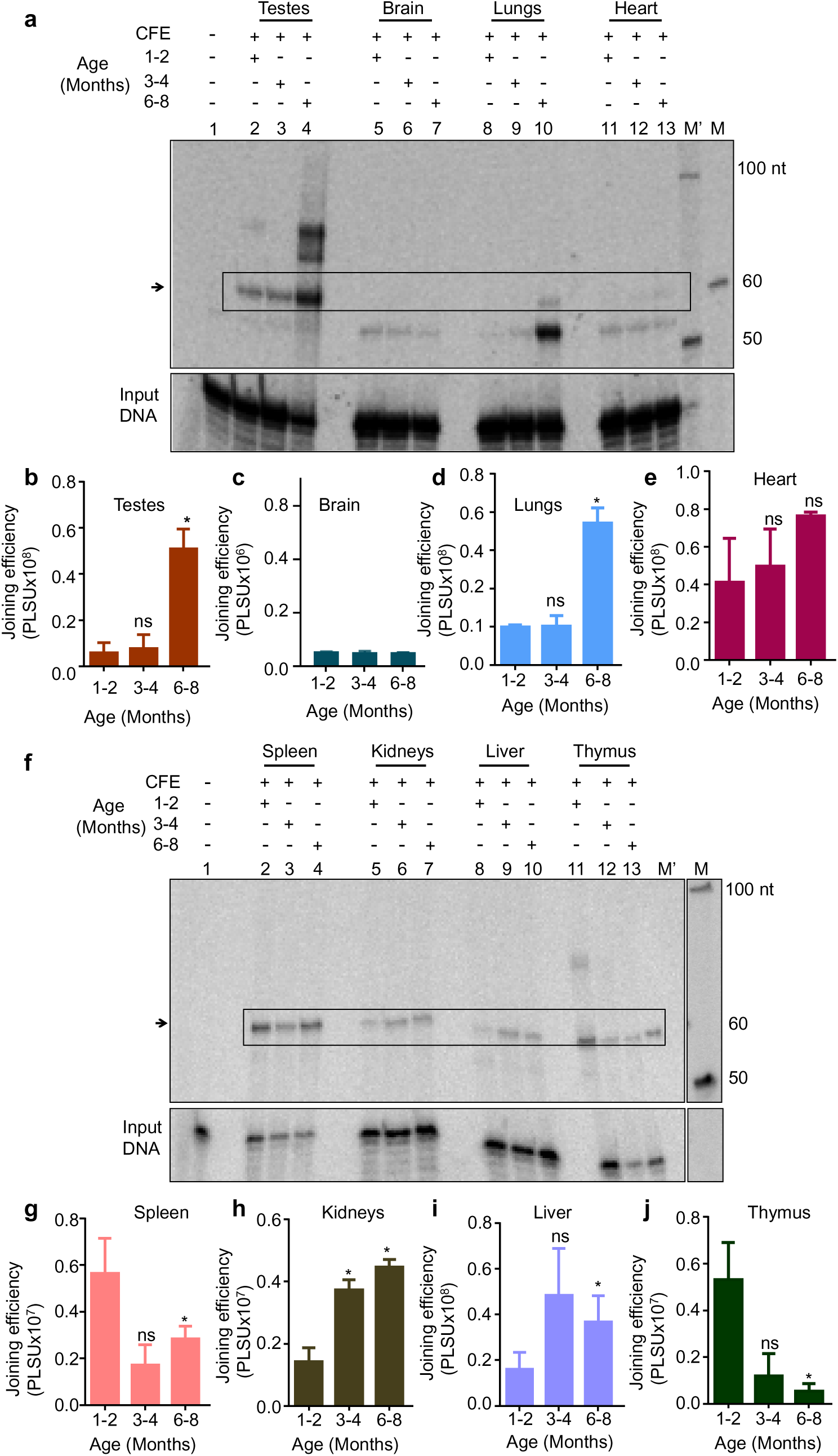
Age-dependent changes in MMEJ efficiency across multiple rat tissues using a 10-nt microhomology substrate. **(a)** Denaturing PAGE showing MMEJ activity in cell-free extracts (CFEs) from early, middle, and late-aged rats. A 10-nt microhomology-flanked substrate was incubated with CFEs from the testes (lanes 2–4), brain (lanes 5–7), lungs (lanes 8–10), and heart (lanes 11–13) in MMEJ buffer (10 mM Tris_⋅_HCl pH 8.0, 20 mM MgCl_₂_, 1 mM ATP, 10% PEG 8000, 1 mM DTT) for 1 hour at 30°C (testes) or 37°C (other tissues). Boxes and arrows indicate MMEJ products. ‘M’ represents the 50-nt marker, and ‘M′’ denotes the 60-nt MMEJ product. Lane 1 is the substrate-only (no-template) control; lane 2 is the no-protein control. **(b–e)** Bar graphs showing quantification of MMEJ activity in CFEs from testes (b), brain (c), lungs (d), and heart **(e)** across early, middle, and late-aged groups. **(f)** Denaturing PAGE showing age-related MMEJ activity in the spleen (lanes 2–4), kidneys (lanes 5–7), liver (lanes 8–10), and thymus (lanes 11–13), with lane 1 as the no-protein control. Reaction conditions were identical to panel **(a)**. Boxes and arrows indicate MMEJ products. **(g–j)** Bar graphs quantifying MMEJ activity in spleen **(g)**, kidneys **(h)**, liver **(i)**, and thymus **(j)** across the three age groups. All quantitative data in panels **(b–e) and (g–j)** represent mean values from at least three independent experiments using extracts from separate animals. Error bars indicate the standard error of the mean (S.E.M.). Statistical significance is denoted as *P < 0.05, **P < 0.01, ***P < 0.001.

In the case of lung extracts, a distinct shift was seen, while MMEJ activity was undetectable in both young and middle-aged groups; a pronounced induction occurred in the aged group (Figure 3a, d; lanes 8–10). This increase in MMEJ activity in the lungs was unlikely to result from differences in CFE protein amounts between age groups, as the protein inputs were normalized across all cohorts (Supplementary Figure S1c), ensuring consistency with previous results (Figure 2h & j). In contrast, only minimal MMEJ was detected in the heart, with a modest but non-significant increase in aged groups (Figure 3a, e; lanes 11–13). These findings reveal a previously unrecognized, age-related activation of MMEJ in select somatic tissues, while others, such as the brain, appear resistant to such changes.

Interestingly, not all tissues followed a pattern of increased MMEJ with age. In both the spleen and thymus, there was a decline in MMEJ efficiency (Figure 3f, lanes 2– 3 and 11–13). The decrease in the middle-aged group was not significant, but in the aged group, the reduction was significant compared to the young cohort (Figure 3g & 3j). These results suggest that MMEJ activity is suppressed with age, particularly in immune-related organs.

In contrast, kidney and liver tissues exhibited an age-associated increase in MMEJ efficiency, with notable upregulation of repair activity observed in these tissues in the aged cohort (Figure 3f). In the kidney, a significant rise in activity was observed in middle-aged animals, which further increased in the aged group, indicating a progressive enhancement of MMEJ efficiency with age (Figure 3f & h, lanes 5–7). In the case of the liver, MMEJ efficiency increased in middle-aged animals, although the change was not statistically significant. However, a significant increase was observed in the aged group (Figure 3f & i, lanes 8–10). These findings reveal a tissue-specific landscape of MMEJ regulation during ageing. While some tissues, such as the testes, lung, heart, kidney, and liver, exhibit upregulation of MMEJ with age, others, like the spleen and thymus, undergo a progressive decline. The consistent absence of MMEJ in the brain further highlights the tightly controlled engagement of this pathway in specific tissues. This pattern underscores the complex interplay between tissue identity, ageing, and DNA repair strategy, with potential implications for age-associated genome maintenance and disease susceptibility.

### Emergence of MMEJ activity in aged lung tissue correlates with Increased FEN1 expression

One of the most striking findings of this study was the age-dependent induction of MMEJ activity in lung tissue—a largely non-dividing organ that exhibited no detectable repair activity in young animals. In contrast to the kidneys, where MMEJ activity increased gradually with age, the lungs displayed a robust and discrete activation of MMEJ exclusively in the aged group (Figure 3), suggesting a distinct regulatory mechanism. To investigate the molecular basis underlying this shift, we analyzed the expression of core MMEJ-associated proteins in lung CFEs across all age groups. While FEN1, Ligase III, and PARP1 levels increased with age, only the upregulation of FEN1 reached statistical significance (Figure 4a, b,c, and e). Importantly, this upregulation strongly correlated with the enhanced MMEJ activity observed in aged lung tissue (Figure 3a).

**Figure 4:**
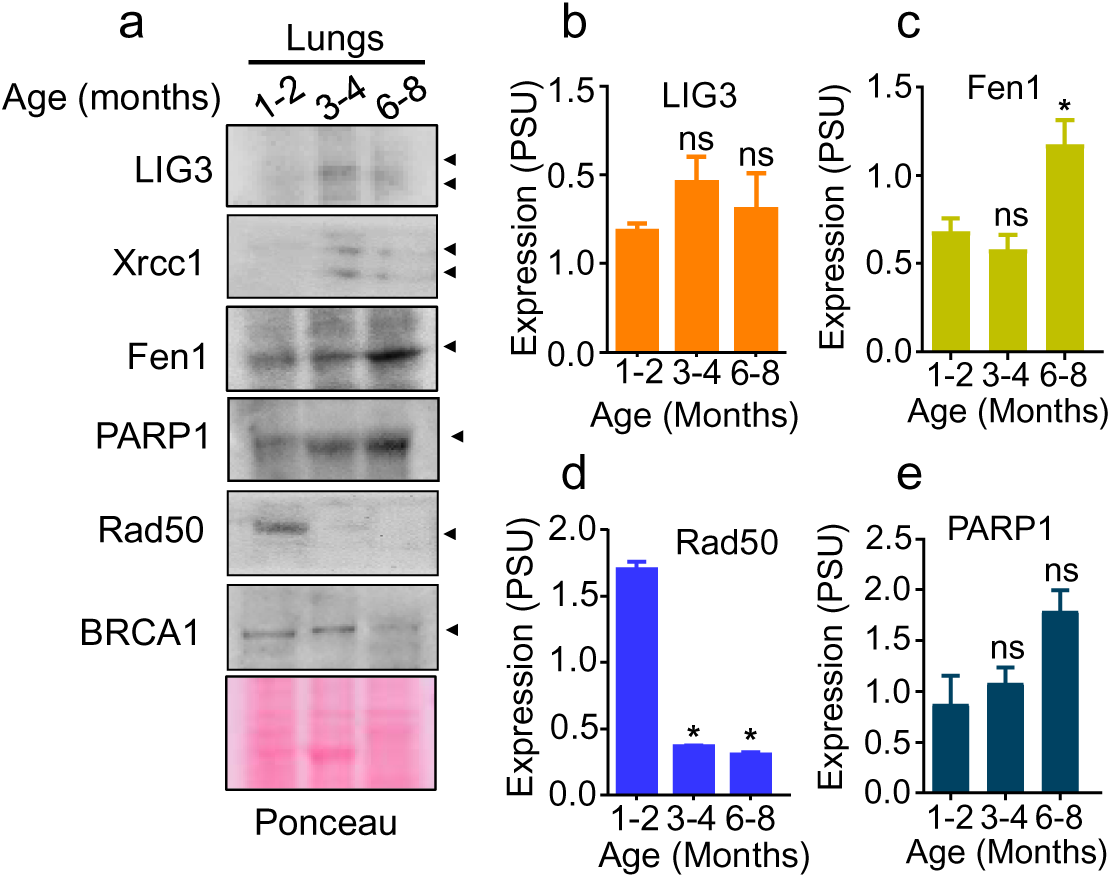
Age-Related Changes in MMEJ Protein Expression in Rat Lungs: Western blots showing the change in expression of LIG3, FEN1, PARP1, and Rad50 in lungs with age. Ponceau served as a loading control. Bar graphs **(b-e)** show the relative change in LIG3, FEN1, Rad50 and PARP1 expression levels in lungs based on levels detected in western blotting. Statistical significance is indicated as *P<0.05; **P<0.01; ***P<0.001, with error bars representing mean ± S.E.M. Expression levels presented in PSU (Protein Signal Units).

In contrast, RAD50 expression declined with age (Figure 4a, d). These data suggest that the age-associated increase in FEN1—potentially in concert with Ligase III and PARP1—may facilitate the activation of MMEJ in ageing lung tissue. This tissue-specific modulation of repair capacity reveals dynamic reprogramming of the DNA damage response during ageing. It provides new insight into genome maintenance mechanisms in post-mitotic organs such as the lung.

Given that ageing induces tissue-dependent shifts in DNA repair pathway utilization, with the lung exhibiting a pronounced reliance on MMEJ in later life, likely driven by changes in the expression levels of MMEJ-associated proteins, especially FEN1. To evaluate whether this trend is conserved in other organs, we extended our analysis to the thymus—a lymphoid tissue known for its high MMEJ activity during early developmental stages (Figure 1c) ((48,54) but shows a marked decline in repair capacity with age (Figure 3f). In aged thymic extracts, compared to early-aged animals, we observed a coordinated downregulation of several core MMEJ-associated proteins, including Ligase III, FEN1, XRCC1, PARP1 and BRCA1 (Supplementary Figure S2). This decrease in protein abundance was consistent with the significant reduction in MMEJ efficiency detected using the 10-nt microhomology substrate (Figure 3f). These findings suggest that the diminished repair capacity in the thymus is driven, at least in part, by reduced expression of essential MMEJ components. Collectively, our data reveal that both the activation and suppression of MMEJ during ageing are governed by tissue-specific regulation of its core protein machinery, reflecting the complex and adaptive nature of genome maintenance pathways across the lifespan.

Given that microhomology length is a critical determinant of MMEJ efficiency and has been directly associated with the frequency of DNA deletion events that accumulate with age (56–58). We next sought to investigate how this parameter modulates MMEJ activity in the context of ageing. To this end, we employed a panel of engineered MMEJ substrates containing microhomology regions of increasing lengths, 13 and 16 nucleotides, with their sequences and design specifications provided in Tables 1 and 2, respectively. This design allowed us to systematically assess the influence of increasing microhomology length on end-joining efficiency in each tissue at different ageing stages. By comparing these patterns across ageing stages, we aimed to uncover how age-related shifts in DNA repair preference and fidelity might contribute to the accumulation of deletion-prone repair events in a tissue-specific manner.

### MMEJ activity with the 13 nt microhomology substrate increased in the testes, brain, and lungs

To investigate the effect of microhomology length on MMEJ efficiency across tissues and ageing stages, we assessed end-joining activity using a 13-nt microhomology substrate. The resulting 65-nt joined product showed a general increase of MMEJ activity, particularly in tissues where repair was weak or undetectable with the 10-nt substrate, indicating that increased microhomology length facilitates more efficient engagement of the MMEJ pathway (Supplementary Figure S3).

In the testes and lungs, elevated MMEJ activity was observed with the 13-nt substrate compared to the 10-nt, particularly in the aged group (Supplementary Figure S3a, b, and d), reflecting enhanced substrate utilization under age-associated stress. This suggests that longer microhomology may facilitate more efficient end-joining in these tissues during ageing. Similarly, in brain tissue, where no MMEJ activity was detected with the 10-nt substrate at any age, low levels of repair became detectable with the 13-nt substrate in both middle-aged and aged samples (Supplementary Figure S3a, c). This finding implies that longer microhomology can partially alleviate the suppression of MMEJ in post-mitotic tissues, allowing for some end-joining activity even in non-dividing cells.

In contrast, heart tissue exhibited consistently low MMEJ activity across all age groups, showing no significant age-related changes regardless of the microhomology length used (10-nt or 13-nt substrates) (Supplementary Figure S3a, e). However, the joining intensity appeared slightly enhanced with the 13-nt substrate compared to the 10-nt, suggesting a modest sensitivity to microhomology length despite the overall low repair capacity in this tissue.

### Age-associated modulation of MMEJ activity across tissues using a 16-nt microhomology substrate

To deepen our understanding of how microhomology length influences end joining with age, we expanded our analysis to include a 16-nt microhomology substrate, which generates a 68-nt joined product. This longer substrate allowed us to assess the impact of extended microhomologies on MMEJ efficiency across various tissues and age groups. By incorporating this substrate, we sought to assess whether ageing alters the utilization of more extended microhomology sequences in end-joining processes, which could potentially reveal tissue-specific changes in repair dynamics and highlight subtle age-related shifts in repair pathway preference. In the testes, end joining continued to increase with age (Figure 5a, lanes 2–4), consistent with trends observed with the shorter 10- and 13-nt substrates (Figures 2 and supplementary figure 3). However, the age-related increase seen with the 16-nt substrate was more modest and did not reach statistical significance (Figures 5a&b).

**Figure 5:**
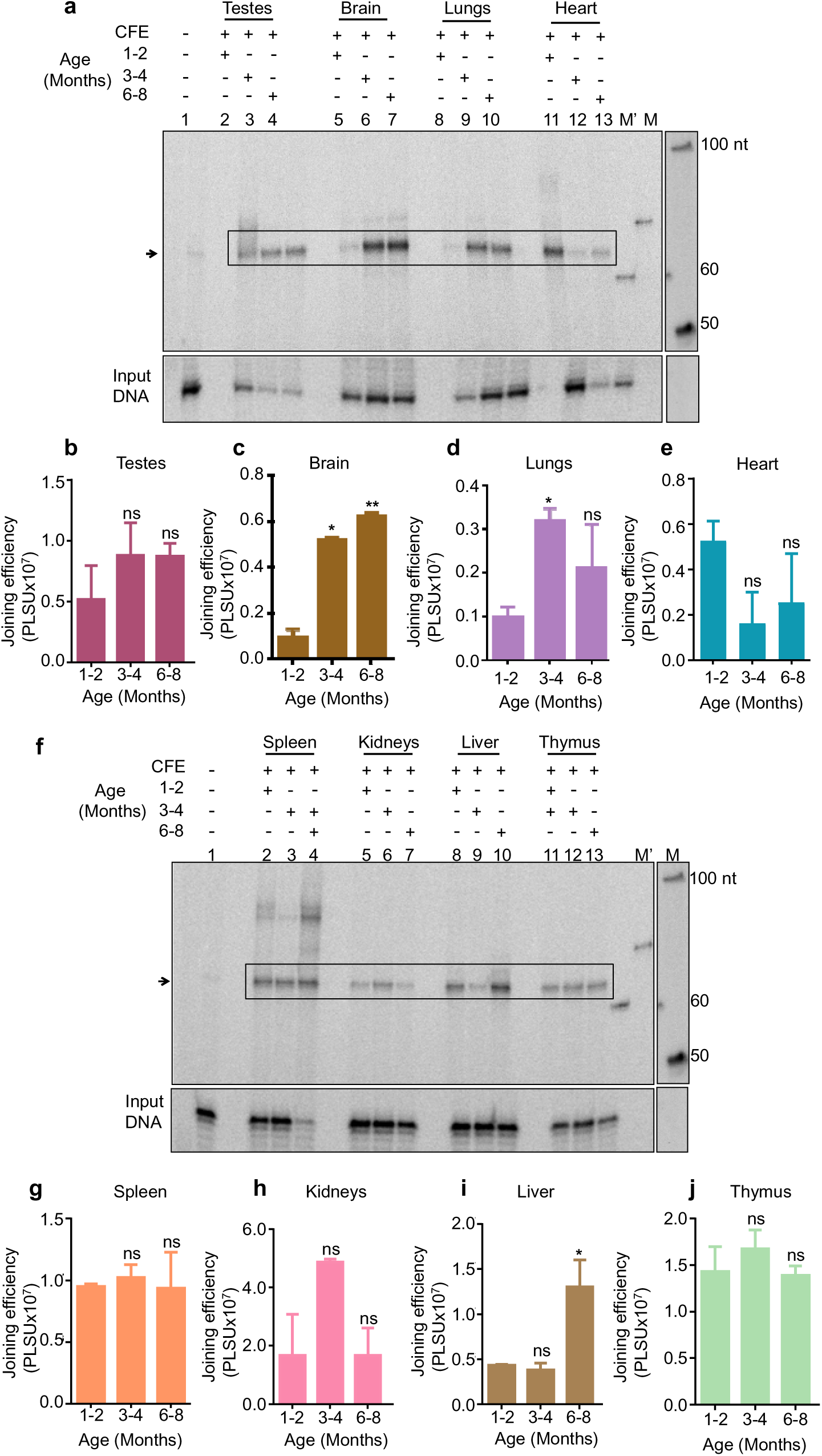
Age-dependent changes in MMEJ efficiency in the testes, brain, lungs, and heart of rats using a 16 nt microhomology substrate. Panel **(a)** shows changes in MMEJ efficiency with age across various organs. A 16 nt microhomology substrate was incubated with CFE from early, middle, and late-aged rats from the testes (lanes 2-4), brain (lanes 5-7), lungs (lanes 8-10), and heart (lanes 11-13) in a buffer containing 10 mM Tris⋅HCl (pH 8.0), 20 mM MgCl_₂_, 1 mM ATP, 10% PEG 8000, and 1 mM DTT for 1 h at 37°C. Joining products were resolved on a 10% denaturing PAGE. Boxes and arrows indicate homology-dependent end-joining products. M represents the 50 nt ladder, M’ means the 60 nt markers and lane 1 is the no-protein control. Panel **(b)** presents a bar graph showing changes in joining efficiency in CFE from testes of early, middle, and late-aged rats. Panel **(c)** shows changes in joining efficiency in CFE from the brain. Panel **(d)** presents a bar graph of changes in joining efficiency in CFE from the lungs. Panel **(e)** shows changes in joining efficiency in CFE from the heart. **(f)** shows changes in MMEJ efficiency with age across the spleen, kidneys, liver and thymus organs. A 16 nt microhomology substrate was incubated with CFE from early, middle, and late-aged rats from the spleen (lanes 2-4), kidneys (lanes 5-7), liver (lanes 8-10), and thymus (lanes 11-13) Joining products were resolved on a 10% denaturing PAGE. Boxes and arrows indicate MMEJ products. M represents the 50 nt ladder, M’ represents the 60 nt marker, and lane 1 is the no-protein control. Panel **(g)** presents a bar graph showing changes in joining efficiency in CFE from the spleen, and panel **(h)** shows changes in joining efficiency in CFE from the kidneys. Panel **(i)** presents a bar graph of changes in joining efficiency in CFE from the liver. Panel **(j)** shows changes in joining efficiency in CFE from the thymus, with results from at least three independent experiments and joining efficiency labeled in PSLU. Statistical significance is indicated as *P<0.05; **P<0.01; ***P<0.001, with error bars representing S.E.M.

In brain tissue, no detectable end-joining activity was observed with the 10-nt substrate at any age, and faint repair activity emerged with the 13-nt substrate in middle-aged rats, though it remained weak. However, with the 16-nt substrate, a significant increase in end-joining activity was observed, first emerging in middle-aged brains and becoming more pronounced in the late-aged group (Figure 5a & c, lanes 5– 7). This shift suggests that, with advancing age, the brain may increasingly rely on joining pathways utilizing longer microhomologies, likely in response to accumulated DNA damage. As a post-mitotic tissue, this enhanced engagement of microhomology-dependent end joining may represent an adaptive strategy to sustain genomic integrity over time.

The influence of microhomology length on end-joining activity was particularly evident in the lungs, where repair dynamics showed a distinct age-dependent pattern. With the 16-nt substrate, end-joining activity was minimal in young rats, peaked in the middle-aged group, and then declined in late age (Figure 5a & d, lanes 8–10). In contrast, the 10- and 13-nt substrates showed only minimal activity in young rats, with significant repair observed primarily in the late-aged group. These findings suggest that longer microhomologies may enable more efficient repair earlier in the ageing process, potentially providing a compensatory mechanism in response to accumulating DNA damage.

In the heart, we observed a moderate decline in end-joining efficiency in the middle- and late-aged groups compared to the early-age group (Figure 5a, lanes 11– 13). However, this reduction was not statistically significant (Figure 5e). Unlike other tissues, the heart exhibited no significant age-related changes even when assessed with the 10- and 13-nt substrates. These findings suggest that microhomology-dependent end joining in the heart may be less sensitive to age or that cardiac tissue may preferentially rely on alternative DSB repair pathways to preserve genomic stability over time.

End-joining efficiency remained relatively stable across all age groups in the spleen and thymus when using the 16-nt microhomology substrate (Figure 5f & g, lanes 2–4; Figure 5f & j, lanes 11–13). This consistency suggests that end-joining activity in these tissues is less affected by ageing when longer microhomologies are available, possibly reflecting a more robust or age-resilient repair mechanism. However, when the 10-nt substrate was used, a significant decline in joining efficiency was observed with age, indicating that shorter microhomologies may expose an age-related vulnerability in the repair pathways of these lymphoid tissues. These findings highlight a substrate-length-dependent sensitivity to ageing, where longer microhomologies may buffer against the decline in DNA repair efficiency.

These findings highlight the complex, tissue and substrate-length-dependent regulation of MMEJ during ageing. While some tissues, such as the testes and lungs, showed increased or dynamic shifts in end joining with age, others, like the heart, remained unchanged, suggesting differential reliance on MMEJ pathways. The brain exhibited an increase in repair efficiency only with longer microhomology, pointing to a possible shift in repair preference in response to ageing-related genomic stress. In contrast, lymphoid organs such as the spleen and thymus maintained stable end-joining activity with the 16-nt substrate but significantly declined when shorter microhomologies were used. These observations suggest that ageing influences the end join capacaity in a substrate-dependent manner, where longer microhomology may help buffer age-related declines in repair efficiency in some tissues. In contrast, others maintain stable activity through alternative or more resilient mechanisms. Overall, our data reveal that the ageing process reshapes the engagement and efficiency of MMEJ in a highly tissue-specific and microhomology-sensitive manner.

## Discussion

Genomic instability is a hallmark of ageing, contributing to a wide range of age-associated pathologies, including cancer, neurodegeneration, and immune dysfunction(59). One of the primary mechanisms driving this instability is the age-dependent decline in high-fidelity DSB repair pathways, such as HR and c-NHEJ (27,28,37,38). Our study demonstrates that MMEJ, an error-prone alternative DSB repair pathway, is dynamically reprogrammed across tissues with age. Importantly, MMEJ activity increases in the liver and testes and emerges in kidneys and lungs with age, where it was dormant in the young (Figure 3a and 3f). At the same time, it decreases in immune tissues like the spleen and thymus (Figure 3f). This tissue-specific modulation likely reflects varying proliferative demands and stress responses.

Prior evidence suggests that MMEJ plays a crucial role during mitosis and may compensate for replicative stress when HR and c-NHEJ are compromised (49–51). Our observation of increased MMEJ efficiency with age in proliferative tissues such as the testes and liver supports this model, indicating a potential link between proliferative activity and MMEJ engagement.

In the testes, ageing is associated with elevated proliferation of a dark spermatogonia—reserve stem cells that re-enter the cell cycle in older individuals, potentially heightening DNA replication stress and creating a cellular environment where MMEJ becomes increasingly essential for maintaining genomic stability (60). Although hepatocyte proliferation generally declines with age in the liver, activation of hepatic stellate cells and other regenerative non-parenchymal populations may contribute to a localized increase in proliferation and hence MMEJ (61). This, in turn, could drive a corresponding increase in MMEJ activity as these dividing cells experience replication-associated DNA damage.

Similarly, in the kidneys and lungs—tissues with low or undetectable MMEJ activity in young animals—we observed detectable MMEJ activity with age. This likely reflects age-related shifts in cellular composition and dynamics. Notably, in the ageing lung, although global proliferative capacity declines (62) specific cell types, such as alveolar fibroblasts, pericytes, airway smooth muscle cells, and endothelial cells, retain or even gain proliferative potential (63). These subsets may increasingly rely on MMEJ to manage DNA damage under conditions where canonical repair pathways become less efficient with age.

Together, these findings support a model in which MMEJ is dynamically regulated in response to proliferative demand and cellular stress. Age-associated changes in cell proliferation, particularly in subpopulations within complex tissues, may modulate MMEJ efficiency, enabling tissues to adapt to evolving genomic maintenance needs during ageing.

Interestingly, the increase in MMEJ efficiency observed in the lungs with age aligns with previous studies using an *in vivo* GFP-based assay (42). However, our results diverge somewhat from those of Vaidya et al., who reported a decline in MMEJ activity in the kidneys and an increase in the heart. These discrepancies are likely attributable to differences in experimental design. While Vaidya et al. utilized fibroblasts isolated from individual organs, our study assessed whole tissue extracts, encompassing both proliferating and non-proliferating cell populations. This cellular heterogeneity may profoundly impact DNA repair dynamics and influence the relative engagement of repair pathways. Moreover, Vaidya et al. assessed MMEJ activity using substrates that encompassed a range of microhomology lengths (1–16 nt), without specifying the exact lengths used in each assay. Given the strong dependence of MMEJ efficiency on microhomology length (56), this lack of specificity further limits direct comparability between the two studies.

Despite these differences, our findings are consistent with the broader observation that increased genomic rearrangements and a heightened reliance on MMEJ accompany ageing. In *S. pombe*, age-driven genome instability is characterised by breakpoint junctions containing microhomologies, implicating MMEJ as a major contributor (43). Similarly, in mammalian systems, a shift toward MMEJ has been reported in ageing epithelial cells and in cancer contexts, particularly during epithelial-mesenchymal transition (EMT), where it could be associated with increased mutational burden and resistance to therapy (41). Together, these observations suggest that ageing may influence DNA repair pathway choice, with a possible shift toward more error-prone mechanisms such as MMEJ in specific contexts.

However, this age-associated increase in MMEJ is not universal across all tissues. In contrast to the liver, kidneys, and lungs, where MMEJ activity increased with age, we observed a decline in the spleen and thymus (Figure 3f). The spleen, where MMEJ plays a vital role in antibody class switch recombination in mature B cells, shows reduced activity, likely due to the well-documented age-related decline in B cell populations (64,65). Similarly, the thymus undergoes age-related involution, characterised by the loss of progenitor and epithelial cells, and likely contributes to the observed decrease in MMEJ activity (55,66). Although TCR rearrangement declines with age, it is mediated by classical NHEJ rather than MMEJ (67,68). Therefore, the decrease in MMEJ likely reflects generalized tissue degeneration rather than impaired antigen receptor assembly.

These findings underscore the importance of cellular composition and functional demand in shaping DNA repair dynamics during ageing.

Beyond cell proliferation, additional factors likely contribute to the increased reliance on MMEJ observed with age. Chronic inflammation, a hallmark of ageing, may play a role in this shift, with emerging evidence linking the elevated cGAS-STING pathway to Polθ activity (69). This indicates that an age-associated inflammatory environment can activate MMEJ. Furthermore, as senescent cells accumulate with age (70) they exhibit impaired repair fidelity and may preferentially engage error-prone repair mechanisms such as MMEJ. Epigenetic alterations, including DNA methylation and histone modifications, also regulate DSB repair pathway choice and may facilitate MMEJ activation in ageing tissues (71,72). Hormonal fluctuations, especially in endocrine-sensitive tissues like the testes, may further modulate DNA repair dynamics and increase MMEJ with age (73).

Although previous studies have reported a decline in DSB repair in spermatozoa, they have primarily focused on the HR pathway (74,75). However, our findings reveal an age-associated increase in MMEJ activity in rat testes, suggesting that this error-prone pathway may become more engaged during ageing. In the context of meiosis, emerging evidence indicates that SPO11-induced DSBs, which are essential for initiating homologous recombination, may be subject to repair by MMEJ (76). This shift toward mutagenic end joining in aged germ cells could compromise the fidelity of meiotic recombination, increasing the risk of infertility, miscarriage, and chromosomal abnormalities such as aneuploidy (77,78). While these findings are based on rodent models, they offer critical mechanistic insights into age-related reproductive decline and germline genome instability in mammals, including humans.

Mechanistically, our findings suggest that the functional decline of high-fidelity DNA repair pathways, such as HR and c-NHEJ, may lead to a greater dependence on MMEJ. Under normal conditions, MMEJ is suppressed by core components of HR and c-NHEJ, including BRCA1, Ku70/80, 53BP1, Rad51, and RPA (79–81)(82) The loss or reduction of these essential repair factors with age (29,83,84) creates a more permissive environment for MMEJ activation. While this shift allows cells to adapt to increased DNA damage, it comes at a cost: MMEJ is error-prone and introduces deletions, insertions, and chromosomal rearrangements, exacerbating genomic instability. The interplay of inflammation, epigenetic changes, and hormonal regulation contributes to the increased reliance on MMEJ, further shaping the tissue-specific DNA repair landscape and impacting genome stability in ageing tissues.

Additionally, our findings reveal that ageing modulates the overall efficiency of MMEJ and drives tissue-specific shifts in microhomology preference, reflecting broader reprogramming of DNA repair strategies with age. In highly proliferative tissues such as the testes and liver, MMEJ activity progressively increased with age. This enhancement extended to longer microhomologies in the liver (Figure 5f), whereas in the testes, the increase was more modest and did not reach statistical significance with the longest substrate (Figure 5a). These observations suggest that ageing may augment mutagenic repair capacity in a substrate- and tissue-dependent manner.

Conversely, in lymphoid tissues like the spleen and thymus, we observed an age-dependent decline in MMEJ efficiency with short microhomology substrates (Figure 3f), while repair using longer microhomologies remained relatively preserved (Figure 5f). This selective repression of error-prone repair at shorter repeats may represent a protective adaptation during immune ageing, limiting genomic instability in long-lived or senescent immune cells.

In the brain, a strictly post-mitotic tissue, MMEJ activity was undetectable with 10- and 13-nt substrates (Figure 3a & supplementary figure S3a) but became detectable with the 16-nt substrate in aged animals (Figure 5a). This shift suggests that shorter microhomologies are insufficient to trigger end joining in neural cells, likely due to stringent regulatory controls that suppress the utilisation of shorter microhomologies. Given that MMEJ preferentially utilises short microhomologies, the preference of longer microhomologous sequences may shift the repair toward single-strand annealing (SSA), a distinct pathway that operates on extended homology regions and results in larger deletions(85,86)(87). The emergence of repair activity with the 16-nt substrate likely reflects a transition toward SSA. Importantly, SSA is associated with extensive deletions, and such a shift in repair pathway choice may underlie the accumulation of large genomic rearrangements observed in ageing-related neurodegenerative and neuropsychiatric conditions (57,88–90). However, it is important to acknowledge that the microhomology length distinguishing MMEJ from SSA is not uniformly defined in the literature, and thresholds vary across studies. This lack of consensus highlights the need for more refined criteria and mechanistic clarity in distinguishing between these closely related end-joining pathways.

The lung displayed a particularly striking pattern, with a robust and consistent age-associated increase in MMEJ activity across all tested microhomology lengths (Figure 2f, 3a, supplementary figures 3a, 5a). This broad activation was accompanied by a marked upregulation of FEN1 expression (Figure 4), a critical end-processing nuclease essential for MMEJ. The correlation between increased FEN1 levels and enhanced MMEJ activity suggests a mechanistic link whereby FEN1 upregulation facilitates the age-dependent activation of this error-prone repair pathway in lung tissue. Notably, FEN1 has been previously reported to be overexpressed in lung cancers, particularly in aggressive subtypes (91). Beyond its role in promoting tumour progression and poor prognosis, elevated FEN1 expression has also been implicated in the development of chemoresistance, owing to its ability to sustain mutagenic repair under conditions of replicative stress and genotoxic challenge (92).

These findings raise the possibility that age-driven upregulation of FEN1 may contribute to the increased incidence and treatment resistance of lung cancer in the elderly by enabling a mutagenic repair environment that fosters both genome instability and therapeutic escape. Moreover, the current study also reveals concurrent age-related increases in PARP1 and Ligase III, two other central mediators of the MMEJ pathway (Figure 4). Their coordinated upregulation suggests a broader reprogramming of DNA repair pathway usage with age, shifting from high-fidelity mechanisms toward more error-prone alternatives such as MMEJ. This shift may reflect a compensatory adaptation to accumulated DNA damage in aged, post-mitotic tissues like the lung, but it also creates a vulnerability that cancer cells could exploit.

Since FEN1, PARP1, and Ligase III are all druggable targets, their combined inhibition represents a promising therapeutic strategy. Previous studies have demonstrated that targeting FEN1 can sensitise cancer cells to DNA-damaging agents (91,93). Similarly, PARP1 plays a central role in MMEJ, where it facilitates DNA end processing and recruitment of Ligase III-XRCC1 (94). Additionally, PARP1 inhibitors are already in clinical use for tumours with homologous recombination deficiency. In HR-deficient cells, its inhibition leads to synthetic lethality by trapping PARP1 on DNA and stalling replication forks (95). Our findings suggest that, in the context of cancer, particularly in the lung, a multi-targeted approach to suppress MMEJ could limit tumorigenesis and overcome drug resistance by restoring dependence on more faithful repair mechanisms or inducing synthetic lethality. Future studies should investigate the therapeutic potential of combining FEN1 inhibitors with PARP1 or Ligase III blockade in lung cancer models, particularly in aged systems, to assess whether this strategy can selectively impair tumour growth without compromising genome maintenance in normal tissues. This approach could provide a novel therapeutic avenue for combating age-related lung cancer progression and overcoming the challenges posed by DNA repair pathway rewiring in the ageing population.

This tissue-specific modulation of MMEJ is also relevant in the context of familial cancer-predisposing mutations in genes such as *BRCA1*, *BRCA2*, and *APC*, which disrupt high-fidelity repair pathways like HR and c-NHEJ (96,97). In individuals carrying such mutations, an age-related shift toward MMEJ may further amplify the mutation burden in susceptible tissues, contributing to their elevated cancer risk. By linking ageing, DNA repair pathway reprogramming, and tissue-specific cancer susceptibility, this study provides new insight into how the interplay between intrinsic ageing processes and inherited repair deficiencies shapes disease risk across the lifespan.

Our study demonstrates that ageing drives a tissue-specific shift in DNA repair pathway usage, characterised by increased reliance on the error-prone MMEJ pathway. This shift is evident in proliferative and post-mitotic tissues such as the testes, liver, lungs, and kidneys(Figure 6). It is accompanied by altered microhomology usage and regulation of MMEJ-related factors. In the lungs, elevated MMEJ activity correlates with a significant upregulation of FEN1.

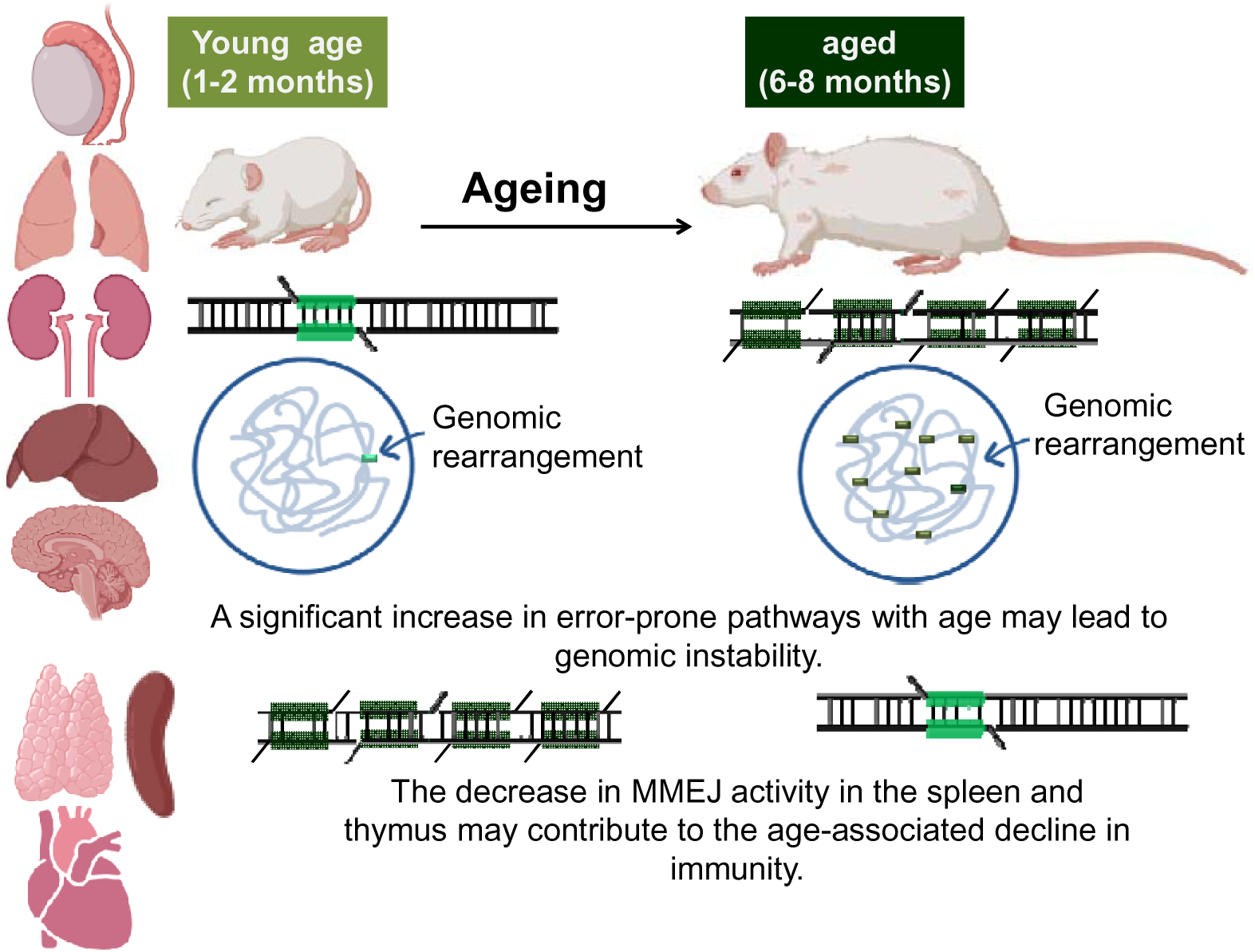

These findings link age-related genomic instability to increased cancer risk and therapy resistance, especially in the lung, and highlight the therapeutic potential of targeting MMEJ components. Moreover, the interaction between ageing, DNA repair reprogramming, and inherited repair deficiencies may underlie tissue-specific cancer susceptibility and age-associated reproductive health decline.

## Supporting information

supplementary infromation

## Abbreviations

APC: Adenomatous Polyposis Coli
BER: Base Excision Repai
BRCA1/2: Breast Cancer Type 1/2 Susceptibility Protein
cGAS: Cyclic GMP-AMP Synthase
c-NHEJ: Classical Nonhomologous End Joining
DNA-PK: DNA-Dependent Protein Kinase
DSB: Double-Strand Break
EMT: Epithelial-Mesenchymal Transition
FEN1: Flap Endonuclease 1
HR: Homologous Recombination
LigIII: DNA Ligase III
MMEJ: Microhomology-Mediated End Joining
MMR: Mismatch Repair
NER: Nucleotide Excision Repair
PARP1: Poly (ADP-Ribose) Polymerase 1
Polθ: DNA Polymerase Theta
SSA: Single-Strand Annealing
SPO11: Sporulation Protein 11
STING: Stimulator of Interferon Genes
TCR: T Cell Receptor
XRCC1: X-ray Repair Cross-Complementing Protein 1
HPRT: GuanineHypoxanthine-Guanine Phosphoribosyltransferase.

## Ethical Approval No

(CAF/Ethics/026/2023).

## Data Availability

All data supporting the findings of this study are included within the article and its supplementary materials.

## Author’s Contributions

The author was solely responsible for the conception, design, data collection, analysis, and writing of this manuscript.

## Consent to Participate

Not applicable

## Consent for Publication

Not applicable.

## Acknowledgements

The author sincerely thanks Prof. Sathees C. Raghavan for his guidance, providing reagents and laboratory space, which were instrumental in completing this study. This work was supported by the Council of Scientific and Industrial Research (CSIR), the Grant-in-Aid for Research Program (GARP), the Department of Biotechnology (DBT), and the Indian Institute of Science (IISc), India.

