## supplementary infromation for "Tissue-Specific Activation of Microhomology-Mediated End Joining with Age Reflects Dynamic Rewiring of DNA Repair in Rats"

**Supplementary information**

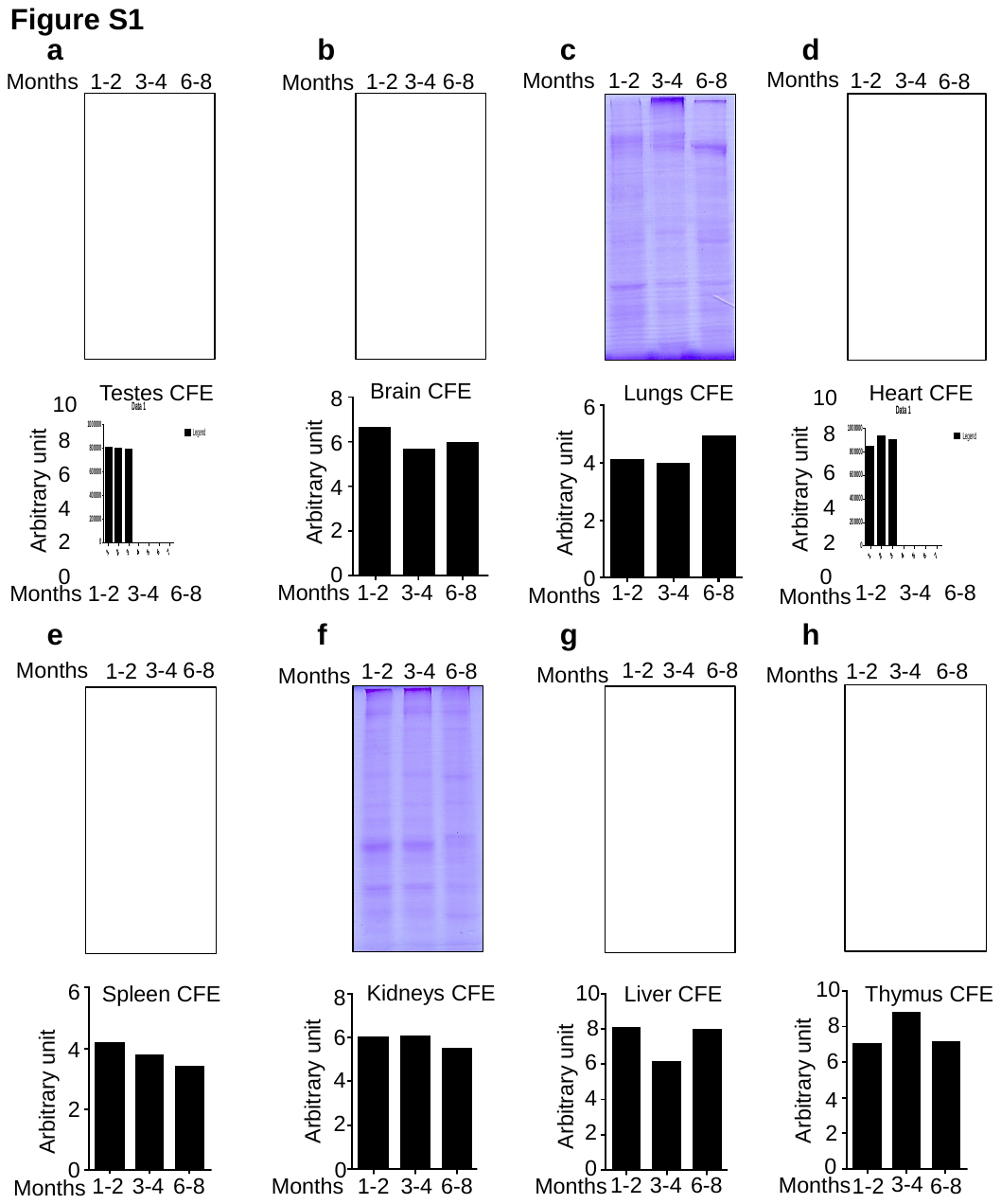

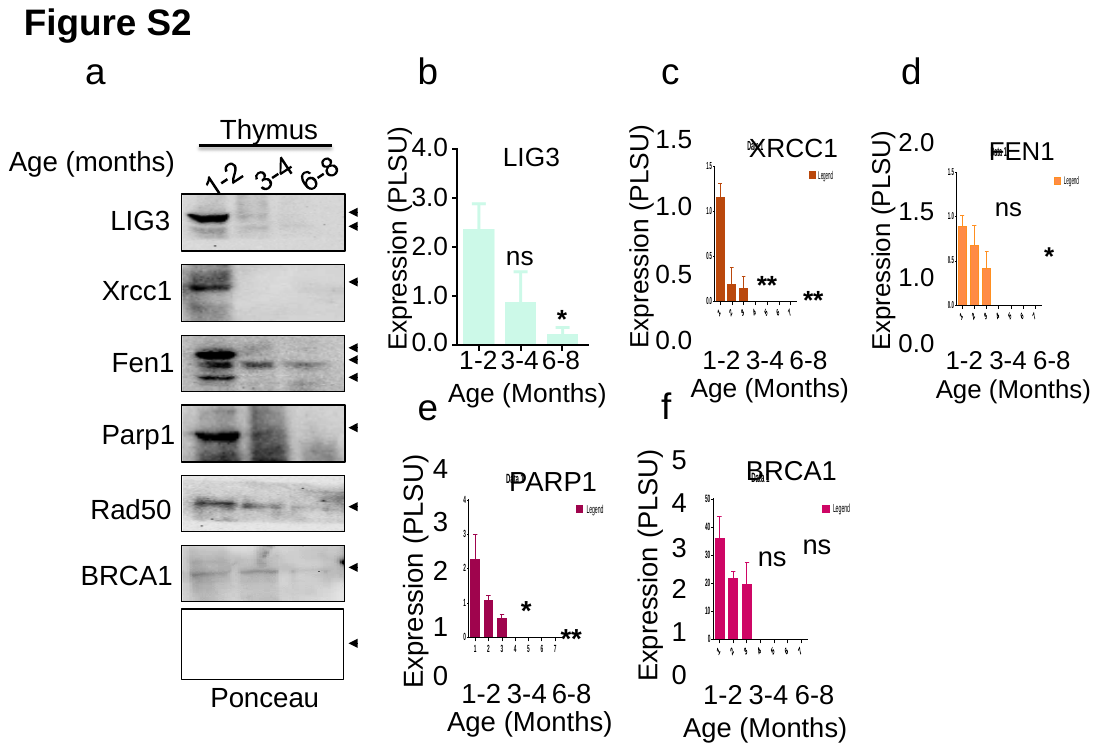

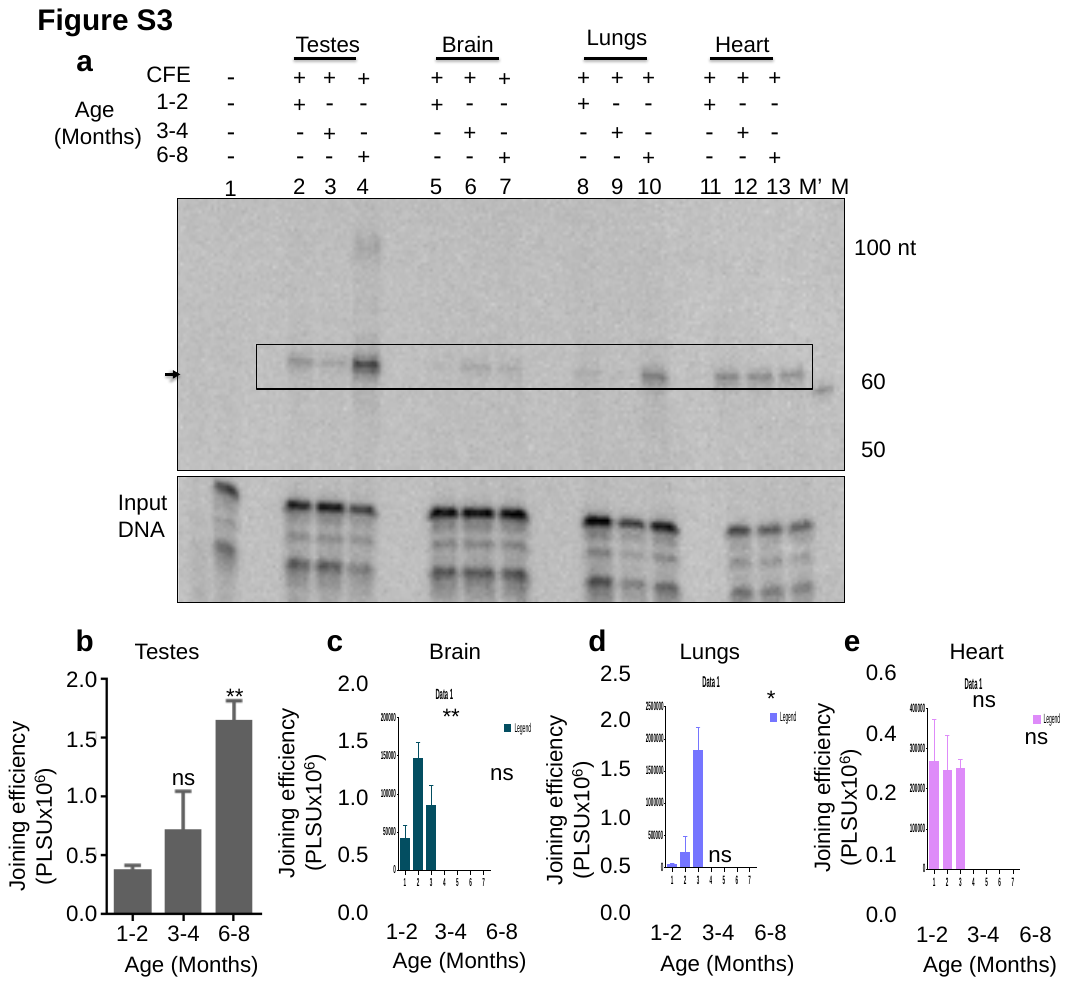

**Figure legends for the supplementary figures**

**Supplementary Figure S1 shows the normalization of CFE across age groups from various rat organs**. Panel **(a)** presents an SDS-PAGE analysis of testis CFE normalisation from early, middle, and late age. Panel **(b)** displays SDS-PAGE analysis of brain CFE normalization across the same age groups. Panel **(c)** illustrates the normalization of lung CFE from early, middle, and late age. Panel **(d)** shows SDS-PAGE analysis for heart CFE normalization across the age groups, while panel **(e)** demonstrates SDS-PAGE analysis for spleen CFE normalization at different ages. Panel **(f)** shows SDS-PAGE analysis of kidneys CFE from early, middle, and late age. Panel **(g)** presents an SDS-PAGE analysis of liver CFE normalization across the same age groups. Panel **(h)** illustrates the normalization of thymus CFE from early, middle, and late age.

**Supplementary Figure S2: Age-related dynamics of MMEJ protein expression in the Thymus:** Western blots showing the change in LIG3, XRCC1, FEN1, BRCA1, and PARP1 in the thymus with age. Ponceau served as a loading control. Bar graphs **(b-f)** show the relative expression levels of LIG3, XRCC1, FEN1, BRCA1, and PARP1 in the thymus based on the level detected in western blotting. Statistical significance is indicated as *P<0.05; **P<0.01; ***P<0.001, with error bars representing mean ± S.E.M. Panel. Expression levels are presented in PSU (Protein Signal Units).

**Supplementary figure S3**: **Age-dependent changes in MMEJ efficiency in the testes, brain, lungs, and heart of rats using a 13 nt microhomology substrate**. Panel **(a)** shows changes in MMEJ efficiency with age across various organs. A 13 nt microhomology substrate was incubated with CFE from early, middle, and late-aged rats from the testes (lanes 2-4), brain (lanes 5-7), lungs (lanes 8-10), and heart (lanes 11-13) in a buffer containing 10 mM Tris⋅HCl (pH 8.0), 20 mM MgCl₂, 1 mM ATP, 10% PEG 8000, and 1 mM DTT for 1 h at 37°C. Joining products were resolved on a 10% denaturing PAGE. Boxes and arrows indicate end joining products. M represents the 50 nt ladder, M' represents the 60 nt marker, and lane 1 is the no-protein control. Panel **(b)** presents a bar graph showing changes in joining efficiency in CFE from testes of early, middle, and late-aged rats. **(c),** shows the changes in joining efficiency in CFE from the brain. Panel **(d)** displays a bar graph of changes in joining efficiency in CFE from the lungs. Panel **(e)** shows changes in joining efficiency in CFE from the heart, with error bars representing S.E.M., with results from at least three independent experiments and joining efficiency labelled in PSLU. Statistical significance is indicated as *P<0.05; **P<0.01; ***P<0.001, with error bars representing S.E.M. Panel.
